# Molecular Underpinnings and Environmental Drivers of Spontaneous Loss of Heterozygosity in *Drosophila* Intestinal Stem Cells

**DOI:** 10.1101/2022.07.21.500951

**Authors:** Lara Al zouabi, Marine Stefanutti, Nick Riddiford, Natalia Rubanova, Mylène Bohec, Nicolas Servant, Allison Bardin

## Abstract

The genome stability of adult stem cells is of particular importance as these cells maintain long-term self-renewal capacity and can contribute extensively to adult tissues. During development and aging, genome mutation leading to loss of heterozygosity (LOH) can uncover recessive phenotypes and be propagated within tissue compartments. This phenomenon occurs in normal human tissues, and is prevalent in pathological genetic conditions and cancers. While previous studies in yeast have defined distinct DNA repair mechanisms that can promote LOH, the predominant pathways underlying LOH in complex somatic tissues of multicellular organisms arenot well understood. In addition, how environmental triggers such as pathogenic bacterial infection may impact LOH is unclear. Here, we investigate the mechanisms giving rise to LOH in adult intestinal stem cells in *Drosophila*. Our data indicate that infection with the enteric pathogenic bacteria, *Erwinia carotovora carotovora 15* but not *Pseudomonas entomophila* increases LOH frequency. Using whole-genome sequencing of somatic LOH events, we demonstrate that they arise primarily via mitotic recombination. Molecular features of recombination sites and genetic evidence argue against formation via break-induced replication and instead support cross-over events arising from double Holliday junction-based repair. This study provides a mechanistic understanding of mitotic recombination in stem cells *in vivo*, an important mediator of LOH.

## Introduction

Studies over the past 10 years have brought to light the fact that healthy, adult tissues are composed of a patchwork of diverse genomes arising from somatic mutation of stem and progenitor cells [1]. A major focus of this body of work has been on the acquisition of somatic point mutations, which are easily detected by short-read sequencing-based approaches [2–4]. Less attention, however, has focused on loss of heterozygosity (LOH) of alleles, more challenging to detect in bulk sequencing. LOH is frequently due to copy number changes, removing one allele, but can also be copy neutral (cnLOH), alternatively referred to as uniparental disomy where 2 identical alleles are present. While the mechanisms underlying development of cnLOH are not fully understood, chromosome mis-segregation would lead to events affecting whole chromosomes. In contrast, repair of DNA damage mediated by recombination machinery would lead to cnLOH events affecting only chromosome arms or segments of chromosomes.

cnLOH, also known as “mitotic recombination”, was originally identified in classic studies from *Drosophila (*[5] and reviewed in [6]), though it also has important beneficial and detrimental consequences on a wide-variety of human pathologies. For example, it has been shown to underlie the spontaneous cure of skin diseases such as Ichthyosis and Epidermolysis Bullosa [7–10] and hematopoietic pathologies like Diamond-Blackfan anemia [11–14]. In these instances, a heterozygous dominant disease-causing allele is reverted to a wild-type allele upon recombination between homologous chromosomes followed by cell division (reviewed in [15]). Mitotic recombination plays a substantial role in both sporadic and familial cancers, particularly problematic for individuals who have germline mutations in *adenomatous polyposis coli* (*APC*), *retinoblastoma* (*Rb*) and *neurofibromatosis 1* (*NF1*) [16–20]. Futhermore, cnLOH has been proposed to represent an important vulnerability in cancers [21]. A better understanding of the process of mitotic recombination and its impact on healthy tissues and disease pathologies is merited.

Mechanistic insight of the molecular events of mitotic recombination has come largely from studies using budding and fission yeast [22–25]. In response to a DNA break, homologous recombination can proceed either through the classic double-strand break repair pathway using a double Holliday Junction intermediate (dHJ) [26], via synthesis-dependent strand annealing, or by Break-Induced Replication (BIR) pathway (reviewed in [25, 27, 28]). However, only the classic dHJ and BIR pathways lead to large stretches of LOH along chromosome arms. In the classic dHJ pathway, DNA break repair proceeds via dHJ intermediates which are often processed by dissolution, creating non-crossover products [29, 30]. Alternatively dHJs are resolved via cleavage by endonucleases, which can promote crossover leading to LOH of heterozygous alleles [29]. Break-induced replication, on the other hand, involves error-prone synthesis of large portions of chromosomes resulting in LOH of heterozygous alleles within the copied region [25, 28]. Studies in mammalian cell culture and in *Drosophila* germline cells have provided additional insight into molecular mechanisms of mitotic recombination and indicated a large degree of conservation of enzymes and processes [31–37]. In particular, *Drosophila* studies revealed that mitotic recombination is normally suppressed by the activity of DNA pol theta-mediated end joining [34]. Nevertheless, many questions are outstanding. The roles of repair by the classic dHJ pathway versus BIR in somatic tissues of Metazoa, for example, is currently unknown.

The *Drosophila* adult midgut has become a powerful model system for understanding healthy tissue dynamics as well as cancer initiation, tumor progression and aging (reviewed in [38]). The intestinal epithelium is composed of differentiated enterocytes (ECs) and enteroendocrine cells (EEs) that are replaced by the asymmetric divisions of intestinal stem cells (ISCs), which are the primary dividing cell type in the midgut [39, 40]. While tissue turnover is slow in unchallenged intestines, ISC proliferation can be rapidly induced by damaging agents such as pathogenic bacteria [41–43], and other damaging agents [44, 45]. The microbiome also impacts the progression of genetically-induced tumors [46, 47]. Additionally, induced tumor models in the *Drosophila* intestine have defined cell-autonomous and non-cell-autonomous signals from the surrounding cellular niche that promote tumor growth and survival signals [48–57]. Thus, the fly gut has provided insight into tumor progression using induced tumor models.

Previously, we demonstrated that the intestine is prone to spontaneous tumor initiation through somatic mutation of ISCs involving gene deletion, chromosome rearrangement, and transposon insertion [58–60]. Furthermore, our findings suggested that a separate process, likely depending on recombination machinery, can promote LOH [58]. Here, we take advantage of the fly intestine model to address the underlying molecular mechanisms and drivers of spontaneously arising LOH in adult stem cells.

## Results

### Spontaneous loss of heterozygosity increases with age

In order to systematically study mechanisms and frequencies of LOH, we chose a null allele of *Suppressor of Hairless* (*Su(H)*) on Chromosome (Chr) 2L (*Su(H)^Δ47^, “Su(H)-”)* as a marker gene. It encodes a transcriptional factor of the Notch pathway and its loss-of-function leads to a readily detectable tumor phenotype phenocopying that of *Notch* with large clones composed of neoplastic ISCs and EE cells [61]. The midguts of a majority of aged *Su(H)-*/+ female flies presented an overall wild-type appearance, composed of large polyploid enterocytes (ECs) with interspersed enteroendocrine (EE cells) and diploid progenitor cells (ISCs and EBs; **Figure 1A-1C**). Less frequently, aged *Su(H)-*/+ midguts were detected with patches of tissue with a *Su(H)* loss-of-function phenotype comprised of an accumulation of Delta (Dl) positive ISCs and Prospero+ (Pros+) EEs were apparent (**Figure 1D, 1E**). These data suggest that, as we previously demonstrated for other Notch pathway components [58], the spontaneous inactivation of the wild-type allele of *Su(H)* occurs during aging representing LOH events.

**Figure 1:**
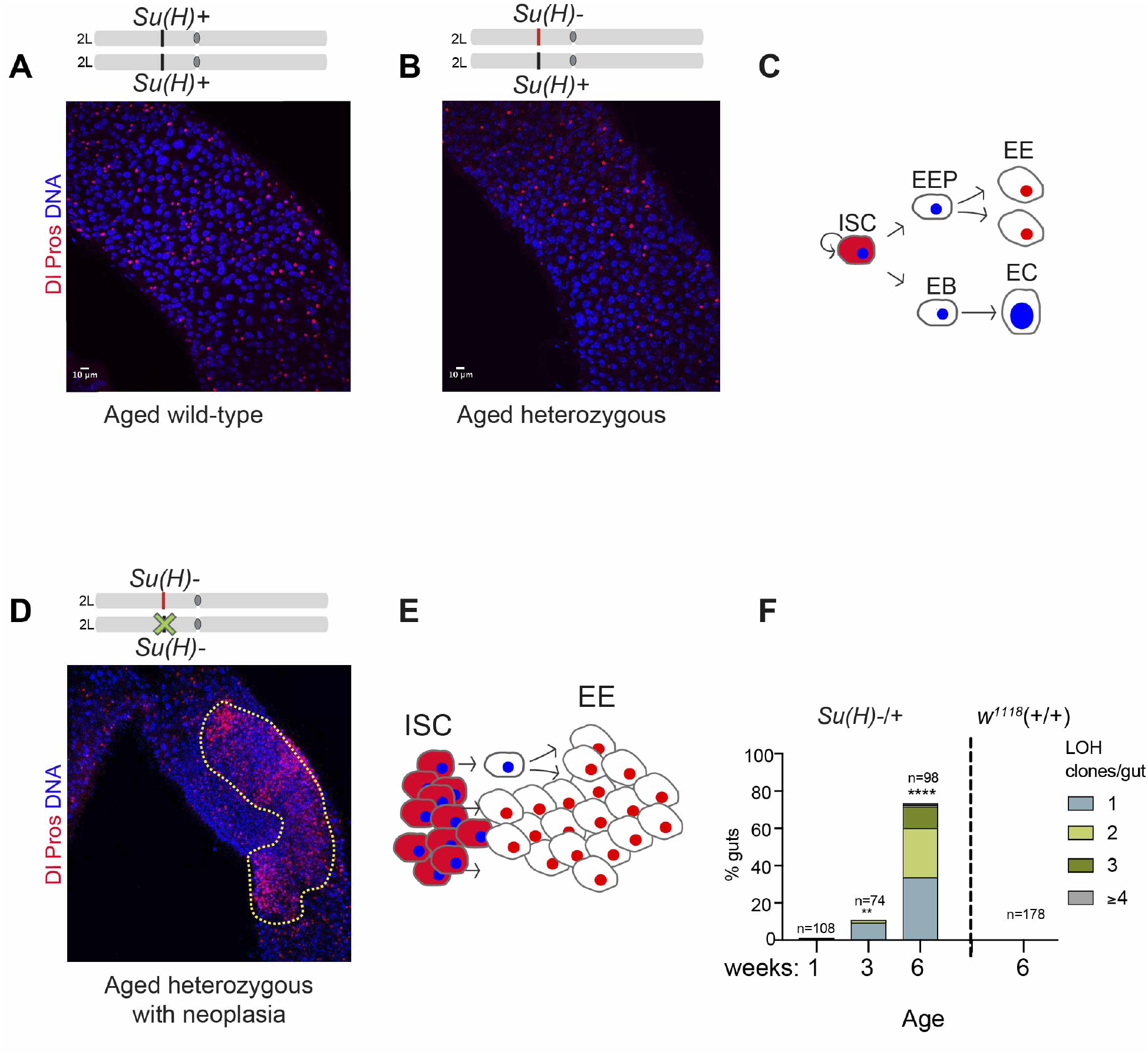
Aging ISCs acquire spontaneous LOH events. **(A)** A wild-type aged gut at 6 weeks. Polyploid enterocytes (ECs), identified by the large nuclear size (DAPI, in BLUE), are the primary cell type in the gut and are interspersed with diploid enteroendocrine cells (EEs, Pros, nuclear RED), and ISCs (cytoplasmic puncta of Delta [Dl] staining RED). **(B)** A heterozygous aged gut at 6 weeks. A majority of tissue in the intestines of Su(H)-/+ flies were like that of wild-type flies. **(C)** ISC lineage: ISCs divide to self-renew and produce EB progenitors that directly differentiate into ECs, representing ∼90% of cells in the tissue. Less frequently, around 10% of ISCs, divide to self-renew and produce EE progenitors, which divide once to make 2 EE cells. **(D)** An example of a heterozygous Su(H)-/+ midgut at 6 weeks of age with a neoplastic clone showing loss-of-function phenotype (outlined in yellow), composed of an excess of Dl+ ISCs and Pros+ EEs **(E)**. **(F)** Frequency of LOH clones in Su(H)-/+ midguts at 1, 3 and 6 weeks of ages. The 1 week time point was used to calculate statistical significance. No midgut showed a neoplastic LOH clone in wild-type intestines. **p < 0.01; ****p < 0.0001(Fisher’s exact test, two-tailed).

The frequency of spontaneously arising LOH clones was found to increase with age. While only 0.9% guts had mutant LOH clones in 1-week old *Su(H)-/+* flies, by 3 weeks of age this rose to 10.8%, and further increased to 73% of midguts by 6 weeks of age (**Figure 1F**). 6-week old wild-type *w^1118^* females (+/+) had interspersed ISCs and EEs and lacked aberrant clusters of these cells seen in LOH conditions (**Figure 1A, 1F)**.

### Infection with the pathogenic enteric bacteria Ecc15 increases loss of heterozygosity

The gut is an organ that responds rapidly to changes in the environment, triggering stem cell proliferation in response to epithelial cell death [62]. To test the role of external environmental factors in driving LOH, we first wanted to validate that environmental alteration occurring in adult life could indeed affect LOH frequencies in the midgut. X-ray irradiation is known to induce chromosomal breaks that lead to LOH in developing larvae [5]. Young 1-week old *Su(H)-/+* flies were irradiated with 40 Gy ionizing radiation (IR) and compared with unirradiated flies at 3- and 6-weeks post-IR (**Figure 2A-2C**). While non-IR treated flies contained 10.8% of guts with spontaneously arising LOH events at 3 weeks, IR-treated flies had a significant increase in LOH events, with 78.4% of midguts containing at least one LOH clones at 3 weeks (**Figure 2B**). In addition, most IR-treated midguts had multiple events (61.9%). At 6 weeks, almost 100% of IR-treated flies had very large LOH clones, often occupying almost the entire midgut, strongly suggesting clone fusion (**Figure 2C**). Importantly, these data illustrate that changes to ISCs during adult life can increase LOH frequency in the midgut.

**Figure 2:**
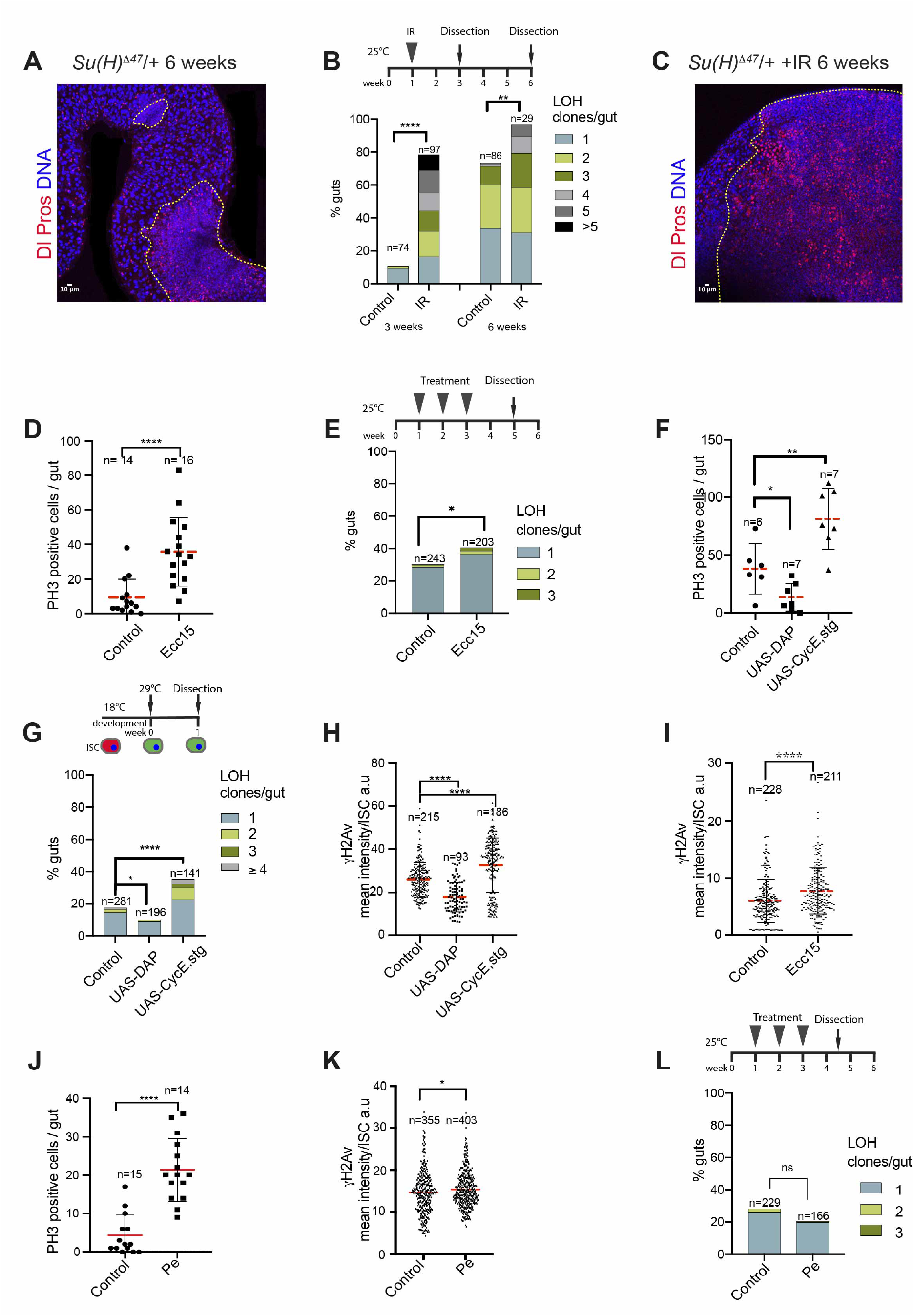
Ecc15 promotes DNA damage and loss of heterozygosity. **(A)** Example of 2 LOH events in aged Su(H)-/+ gut. Dl (ISCs, cytoplasmic RED), Pros (EEs, nuclear RED), DAPI (BLUE). **(B)** A significant increase in LOH frequency was found in Su(H)-/+ flies irradiated at 1-week of age with 40 Gray ionizing radiation (IR), compared to unirradiated flies at 3 and 6 weeks post-IR. Note, control data are also used in Figure 1D. **(C)** An example of a large LOH clone in an aged IR-treated Su(H)-/+ gut outlined in yellow Dl (ISCs, cytoplasmic RED), Pros (EEs, nuclear RED), DAPI (BLUE). **(D)** Infection of 3-5 day old flies with pathogenic bacteria Ecc15 stimulates ISC proliferation 24h after treatment as assessed by Phospho-Histone H3 (PH3), (t-test with Welch’s correction). **(E)** Frequency of LOH clones in Ecc15 treated flies compared with control. Flies were treated for 24 hours in weeks 1, 2 and 3 and were dissected in week 5, providing time for recovery to minimize overall toxicity and avoid lifespan reduction (Fisher’s exact test, two-tailed). **(F)** PH3 quantification of control guts or those overexpressing the cell cycle inhibitor DAP or cell cycle drivers CycE and stg expressed in stem cells using Dl-GAL4 combined with tub-GAL80ts for 7 days (t-test with Welch’s correction). **(G)** Frequency of LOH in midguts of controls and those overexpressing the cell cycle inhibitor DAP or cell cycle drivers CycE and stg expressed in stem cells Dl-GAL4 combined with tub-GAL80ts at 7 days (Fisher’s exact test, two-tailed). **(H)** Comparison of γH2Av mean intensity (arbitrary units) in ISCs overexpressing DAP (n=93, N=5) or cell cycle drivers CycE and stg in ISCs (n=186, N=6) compared with control (n=215, N=8). ‘n’ is the total number of ISCs quantified, ‘N’ is the number of guts assayed. **(I)** Infection of 3-5 day old flies with pathogenic bacteria Ecc15 promoted an increase in γH2Av 24h after treatment (n=211, N=6) compared with control (n=228, N=6; t-test with Welch’s correction). **(J)** 3-5 day old flies dissected immediately after 24 hours of Pe treatment to assess a proliferative response (t-test with Welch’s correction). **(K)** Comparison of γH2Av mean intensity in ISCs in Pe treated (n=403, N=5) and control flies (n=355, N=8) after 24h of treatment of 3-5 days old flies (t-test with Welch’s correction). **(L)** Frequency of LOH clones in Pe treated flies compared with control (Fisher’s exact test, two-tailed). *p < 0.05; **p < 0.01, ****p < 0.0001; not significant (n.s.).

We then wanted to assess whether natural pathogens may impact somatic mutation via LOH in the midgut. The pathogenic bacterial strain *Erwinia carotovora carotovora 15* (*Ecc15*) can infect *Drosophila melanogaster*, cause epithelial damage, and promote ISC proliferation [42]. As previously reported, *Ecc15* treatment led to an increase in mitotic cells after 24 hours of treatment compared to untreated controls (**Figure 2D**). Punctual exposure of *Su(H)-/+* flies to *Ecc15* during 24 hours, once per week for 3 weeks, was performed with midguts assessed at 5 weeks of age. *Ecc15* treatment led to an increase in frequency and number of LOH events compared to controls: 40.6% upon *Ecc15* treatment had at least 1 LOH event compared to 30.3% of guts in the untreated control (**Figure 2E**).

We suspected that the effect of *Ecc15* on LOH might be due to the enhanced number of cells divisions, thereby increasing the likelihood of replicative DNA damage. To test whether increased cell division could alter DNA damage and LOH frequency, we modified cell division rates in adult ISCs (*Dl^GAL4^* combined with *tub-GAL80^ts^*), using either: 1.) the overexpression of the cyclin-dependent kinase inhibitor *Dacapo* (*Dap*), known to block the G1/S transition and thus reduce cell division [63], or 2.) the combined overexpression of *Cyclin E* and *string*, previously shown to increase ISC proliferation [64]. The frequency of proliferation, number of LOH events, and quantity of γH2Av, a mark of DNA damage, were significantly decreased upon *Dap* expression and increased upon combined overexpression of *CycE* and *Stg* (**Figure 2F-H**). Consistent withproliferation driving DNA damage, *Ecc15* treatment was also found to increase γH2Av levels in ISCs (**Figure 2I**). These data suggest that *Ecc15* treatment could potentially impact LOH due to its effect on stem cell proliferation, which may be directly linked to the observed effects on DNA damage.

We then asked whether another pathogenic bacteria resulted in a similar effect on LOH. *Pseudomonas entomophila* (*Pe*) treatment stimulated ISC proliferation as previously reported (**Figure 2J**, [43]), and was found to increase a DNA damage, γH2Av (**Figure 2K**), though to a lesser extent than *Ecc15*. However, treatment with *Pe* did not significantly alter LOH frequency (**Figure 2L**). Together, our findings demonstrate that environmental changes such as gut enteropathogens can influence the frequency of LOH events, but that they are not all equal in their capacity to induce DNA damage or LOH.

### Whole genome sequencing reveals that LOH arises through mitotic recombination

We then wanted to determine the molecular mechanism underlying LOH. While our previous study hinted towards mitotic homologous recombination as a mechanism [58] as LOH frequency was diminished by balancer chromosomes known to suppress recombination, other mechanisms could not definitively be excluded. For example, chromosome loss or deletion might lead to LOH. In addition, LOH occurring via cross-over could not be distinguished from that driven by BIR in our previous study [58]. Finally, a recent study suggested that an unusual chromosome segregation mechanism, “amitosis”, can lead to LOH in the *Drosophila* midgut under conditions of starvation stress [65]. In the proposed mechanism of amitosis, enteroblast progenitor cells of the gut, after 1 round of endoreplication while they are 4n ploidy, are thought to undergo a reductive cell division leading to the segregation of 2 chromosomes originating from the same parent into one daughter cell [65].

Therefore, to differentiate between these mechanisms, whole genome sequencing (WGS) of *Su(H)* LOH clones was performed to determine the molecular nature of inactivation events (**Figure 3A-C**). LOH clones from 5-week-old flies were identified with a GFP reporter of EE cells, which aberrantly accumulate in the tumors as GFP+ clusters (*Su(H)-*/+; *Pros^V1Gal4^*/ *UAS-nlsGFP*; **Figure 3B**). Genomic DNA was then isolated from the “tumor” mutant clone and “normal” head and Illumina paired-end (150bp) sequencing was performed on 28 female and 6 male intestinal neoplastic clones along with the head sample from the same fly as a control of the *Su(H)-*/+; *Pros^V1Gal4^/ UAS-nlsGFP* genetic background **(Supplementary Table 1**, **Figure 3C)**. Four female samples were excluded from further analysis due to low sequencing coverage after mapping to the *Drosophila* genome and removing of duplicate reads (red samples in **Supplementary Table 1**). Importantly, the parental genotypes carried a large number of single nucleotide polymorphisms (SNPs), used in the bioinformatic analysis detailed below.

**Figure 3:**
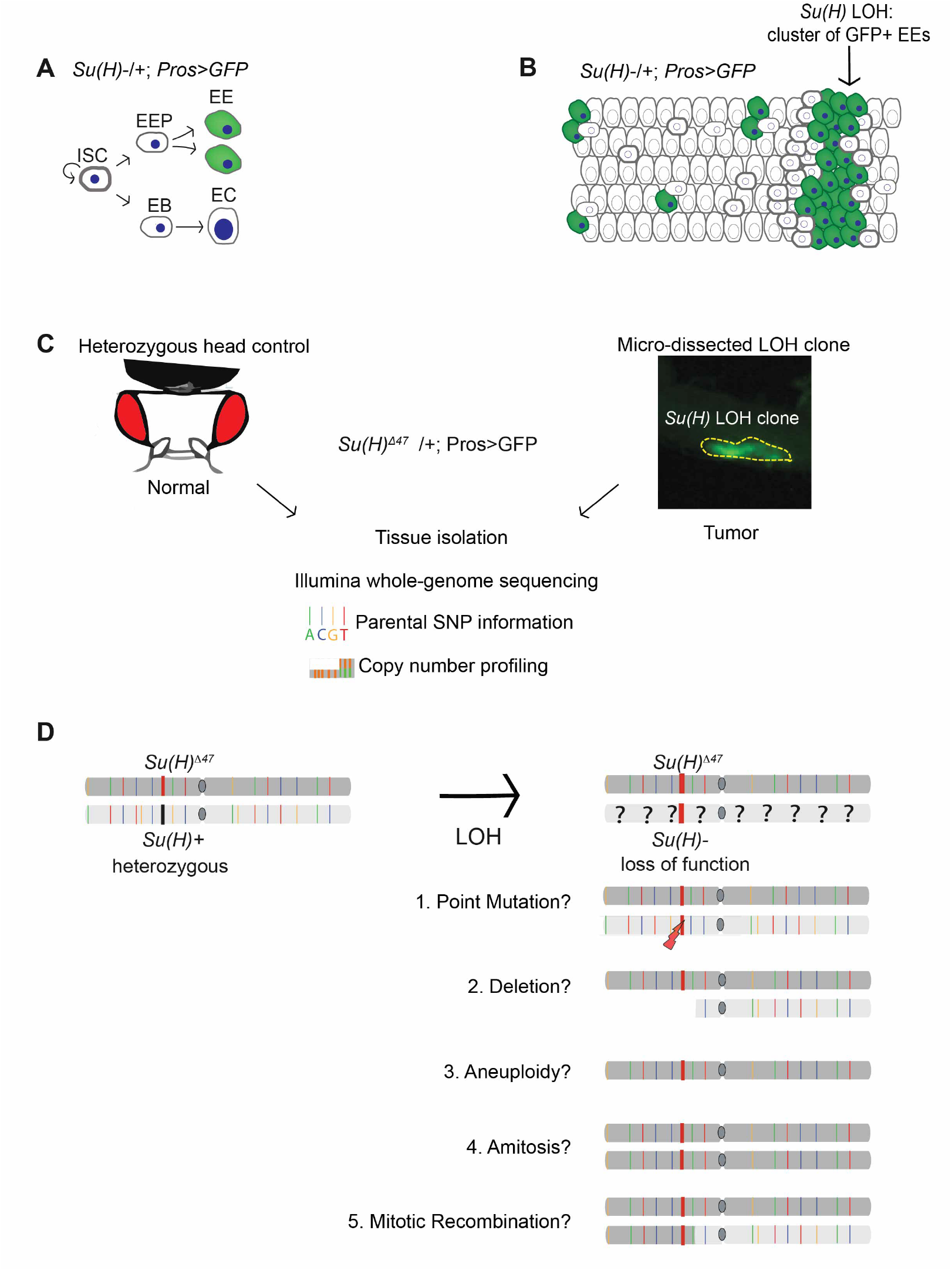
Sequencing setup and analysis predictions. **(A)** ProsGal4 UAS-GFP transgenes drive GFP specifically in EEs allowing for isolation of large green LOH clones composed of ∼1000-5000 cells. **(B)** Schematic of a Su(H) LOH clone recognized by the accumulation (marked by an arrow) of GFP+ EEs. **(C)** The GFP+ LOH clones were microdissected along with the head of the same fly for Illumina paired-end sequencing. Of note, the head was used as a control because it contains primarily diploid cells like the neoplastic LOH (composed of ISCs and EEs) as opposed to adjacent gut tissue, which has complex polyploid genomes due to the presence ECs. Germline single nucleotide polymorphisms (SNPs), somatic point mutations (SNVs), and copy number profiling was assessed. **(D)** SNP, SNV, and copy number profiling distinguish between 5 potential mechanisms by which LOH can arise.

To distinguish between different mechanisms of LOH, whole-genome sequencing (WGS) data of 24 female and 6 male neoplasia samples tumor/normal pairs was analyzed including copy number changes, structural variants, somatic point mutations and changes to the zygosity of parental SNPs (**Figure 3D)**. In the 24 female samples, no point mutations in *Su(H)* or whole chromosomal copy number change occurred Chr 2 on which *Su(H)* is located, ruling out point mutation and aneuploidy as causes of LOH for these samples (**Supplementary Figure 1)**. One sample showed a large structural variant (deletion) spanning 1.38MB of *Su(H)* (sample F20; **Supplementary Figure 2**). In one sample (F28), no LOH or mutation could be found. However, in the majority of samples (22/24 samples), changes in SNP heterozygosity were detected as a shift in zygosity from a variant allele frequency (VAF) of ∼0.5 to 0.75 or above or 0.25 and below on Chr 2L (**Figure 4A**, **4B**, **Supplementary Figure 1)**. As LOH samples have some contaminating wild-type adjacent cells, this number does not go to 1 or 0. The positions of LOH invariably arose at locations between the *Su(H)* locus and the centromere 7.5Mb away and extended throughout the chromosome to the telomere, thereby resulting in LOH of the *Su(H)* locus **(Figure 4A, 4B;** for all 22 samples see **Supplementary Figure 1**). As SNP changes affected only a portion and not the entire chromosome arm, the data supported a mitotic recombination-based mechanism and not amitosis, which would affect the entire chromosome. Further validation of this was obtained taking advantage of the chromosomal position of a UAS-GFP located on Chr 2L (28F3-28F5), more telomeric than the *Su(H)* locus on Chr 2L (35B), whose expression was lost in LOH clones **(Supplementary Figure 1B-C’)**.

**Figure 4:**
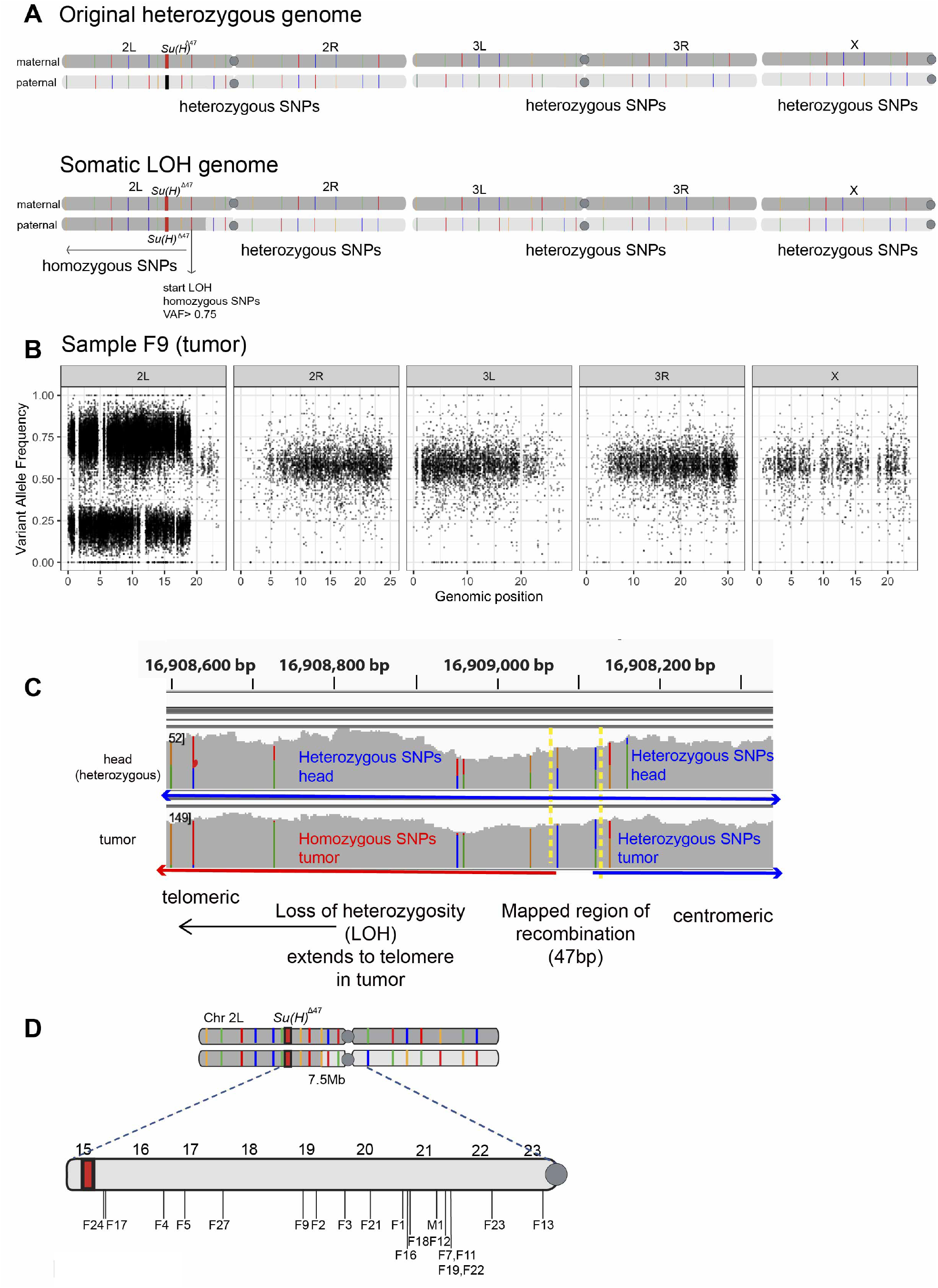
Sequencing Su(H) LOH clones reveals loss of heterozygosity primarily arises though mitotic recombination. **(A)** A schematic corresponding to the Variant Allele Frequency (VAF) plot below. **(B)** A representative VAF plot of a female sample (F9). In this LOH sample, chromosomes 2R, 3L, 3R and X had heterozygous SNPs, represented at ∼0.5 VAF. In contrast, chromosome 2L has undergone an LOH event, with SNPs becoming homozygous (>0.75%) shown in the left panel. In a scenario where the tumor purity is 100%, the VAFs in LOH regions should be 1 (100%) or 0, however given the manual nature of the tumor micro-dissections, contaminating non-tumor cells make up part of the sequenced tumor resulting in deviations from this. **(C)** IGV view of mapping of recombination sites in LOH neoplasias compared to heads. Both head and tumor (LOH) samples show heterozygous SNPs on the right side, seen by coloured SNPs each at roughly 50% of the sequencing depth. In contrast, on the left side, these SNPs now become homozygous in the tumor, where they are still heterozygous in the head. Between the yellow lines is the region where recombination took place, mapped within 47bp the last heterozygous SNP and the first homozygous SNP in tumor. **(D)** All of the mapped recombination sites occurred between Su(H) locus (red) and the centromere in female samples (F1-F27) and the male sample M1. Numbers on the chromosome arm correspond to chromosome coordinates.

Additionally, evidence for LOH via mitotic recombination in 1 of the males samples was also found. A shift in allele frequency of SNPs on Chr 2, resulted in LOH of a large portion of the chromosome arm, including the *Su(H*) locus **(Supplementary Figure 3A-A’)**. The remaining 5 male neoplastic samples had inactivating deletions of the *Notch* locus located on Chr X, hemizygous in males **(Supplementary Figure 3B-B’),** consistent with events that we previously described in wild-type male flies leading to neoplasia [58, 60]. Therefore, in total, mitotic recombination was detected in 23 samples (22 females and 1 male).

These data provide evidence that the primary means of LOH leading to spontaneous neoplasia formation in intestinal stem cells is mitotic recombination. Importantly, our data rule out a major contribution of other mechanisms of LOH including (1) point mutations, (2) deletions, (3) chromosome loss, and (4) amitosis. Furthermore, the long track of LOH extending to the telomere rules out repair mechanisms that result in short tracks of LOH that do not lead to crossover, including gap repair, single-strand annealing, and synthesis-dependent strand annealing [27].

### The Histone Locus Cluster is enriched for sites of mitotic recombination

We next wanted to understand whether there are underlying DNA sequence features that contribute to the DNA damaging event driving mitotic recombination. We were able to map the sites where the recombination occurred in 20 out of the 23 samples, detected as the chromosomal region (sequence) between the first homozygous SNP (from the centromere) and the position of the last heterozygous SNP (from the centromere) in the LOH “tumor” sample (see Methods and **Figure 4C, 4D, Supplementary Table 2).** Recombination sites were mapped from a resolution of 47 bp to 111kb and had a median size of 1865bp (**Supplementary Table 2**). Three samples had low purity of the neoplastic clone precluding mapping of the region of recombination (see Methods). The position of recombination was then compared to various sequence features. There was no significant overlap with mapped R loops [66] and predicted sequences for form non-B-form DNA such as G-quadruplexes, cruciform DNA, though did find an a significant enrichment for short inverted repeats (SIRS, **Supplementary Figure 4**).

Interestingly, 4/20 samples had a recombination site that arose within the *Histone Locus Cluster*, a region of 110 kb within the 7.5 Mb region between the centromere and the *Su(H)* locus (**Figure 5A**). These are only 4/20 samples and, thus, should be interpreted with caution. However, the occurrence of recombination sites within this small region (110 kb) was significantly enriched to what was expected using simulated data (**Figure 5B**). The *Histone Locus Cluster* is an array of 100 copies in tandem each containing 5 Histone genes (H2A, H2B, H3, H4 and H1) (**Figure 5A**), which has features that could contribute to replication problems. First, it has tandem repeats which may cause problems for the replication fork. Secondly, the *Histone Locus Cluster* is also only highly transcribed during S phase when Histones are incorporating into newly synthesized DNA [67], which could possibly make it more prone to replication fork collisions with the transcription machinery and lead to DNA damage. Further investigation of the *Histone Locus Cluster* could provide important insight into genomic features potentially driving mitotic recombination from the homologous chromosome.

**Figure 5:**
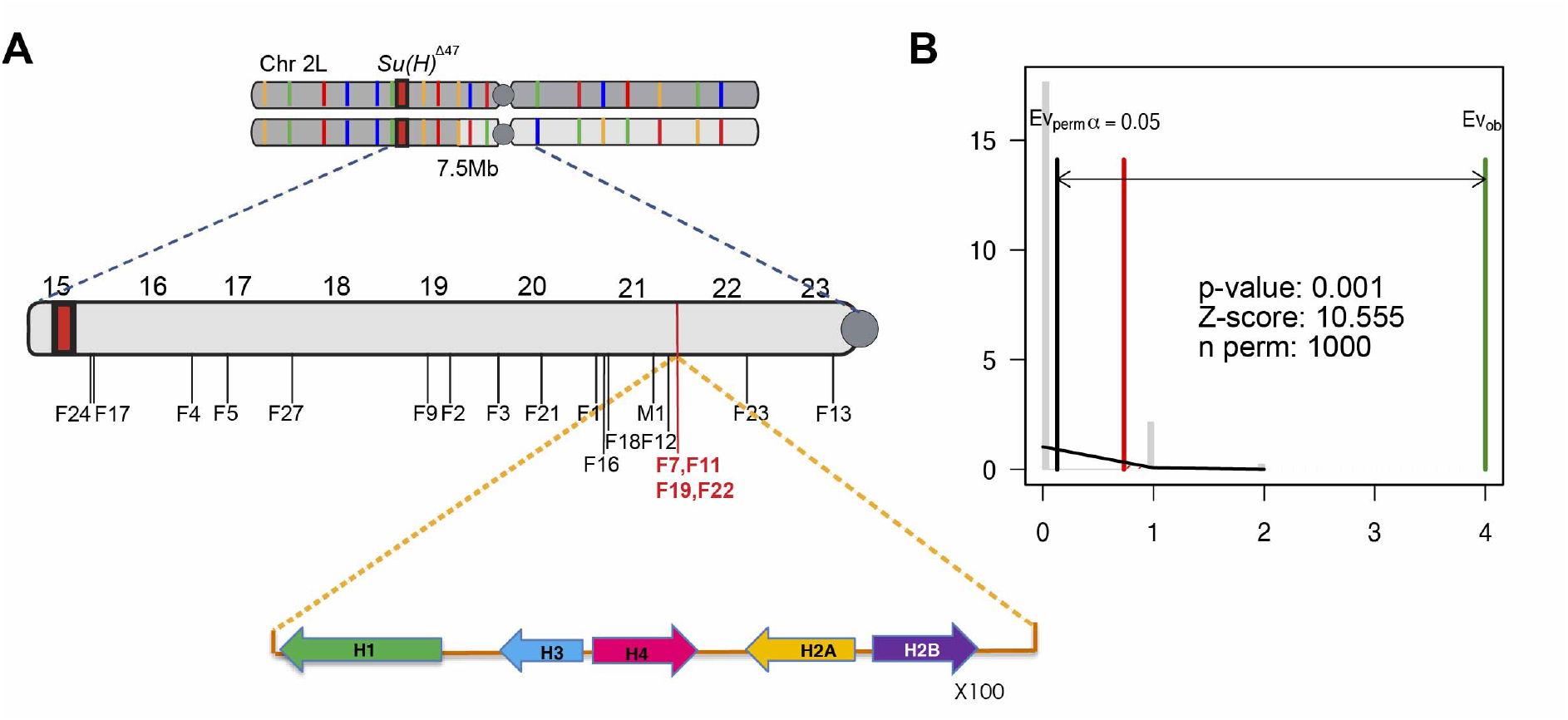
The Histone Locus-Cluster: a putative genomic feature driving mitotic recombination. **(A)** 4 samples (F7, F11, F19 and F22 shown in red) had recombination sites that occurred within the Histone Locus Cluster. Due to the repeat nature, the exact positions could not be mapped within this ∼110kb region. Below the chromosome arm is a schematic of the Histone Locus-Cluster; an array of 100 copies of 5 Histone genes. **(B)** Permutation test carried out using RegioneR, p<0.001. Test association between mapped recombination sites and the Histone Locus-Cluster. Shuffles: 1000 constricted to the region between Su(H) and the centromere (where recombination can be detected).

### Rad51 promotes loss of heterozygosity

Our data above indicate that mitotic recombination plays a major role in LOH in intestinal stem cells. As recombination results from DNA double-strand break repair (DSBR) that utilizes the Rad51 protein to invade a donor DNA molecule **(Figure 6A)** [27, 33], we predicted an involvement of the *Drosophila* Rad51 protein SpnA. We hypothesized that in absence of Spindle-A (SpnA), damaged stem cells would be unable to repair DNA and would consequently die, reducing the number of LOH clones detected. Consistent with this, LOH frequency decreased from 32.6% in *Su(H)-/+* controls to 7.5% in *Su(H)-/+; SpnA^093^/SpnA^057^*null mutants **(Figure 6B)**. No obvious effect on stem cell number occurred in *SpnA^093^/SpnA^057^* mutants, ruling out the trivial explanation that the ISC population is reduced (**Supplementary Figure 5A, B**). Simlar to *SpnA* mutants, there was a reduction in LOH clones upon knockdown of *Rad51 (SpnA)* specifically in adult stem cells (*Dl-Gal4*, *tub-Gal80ts*; **Supplementary Figure 5C,** see Methods for information on scoring). We conclude that LOH arises in ISCs largely from a Rad51-dependent repair mechanism driving mitotic recombination.

**Figure 6:**
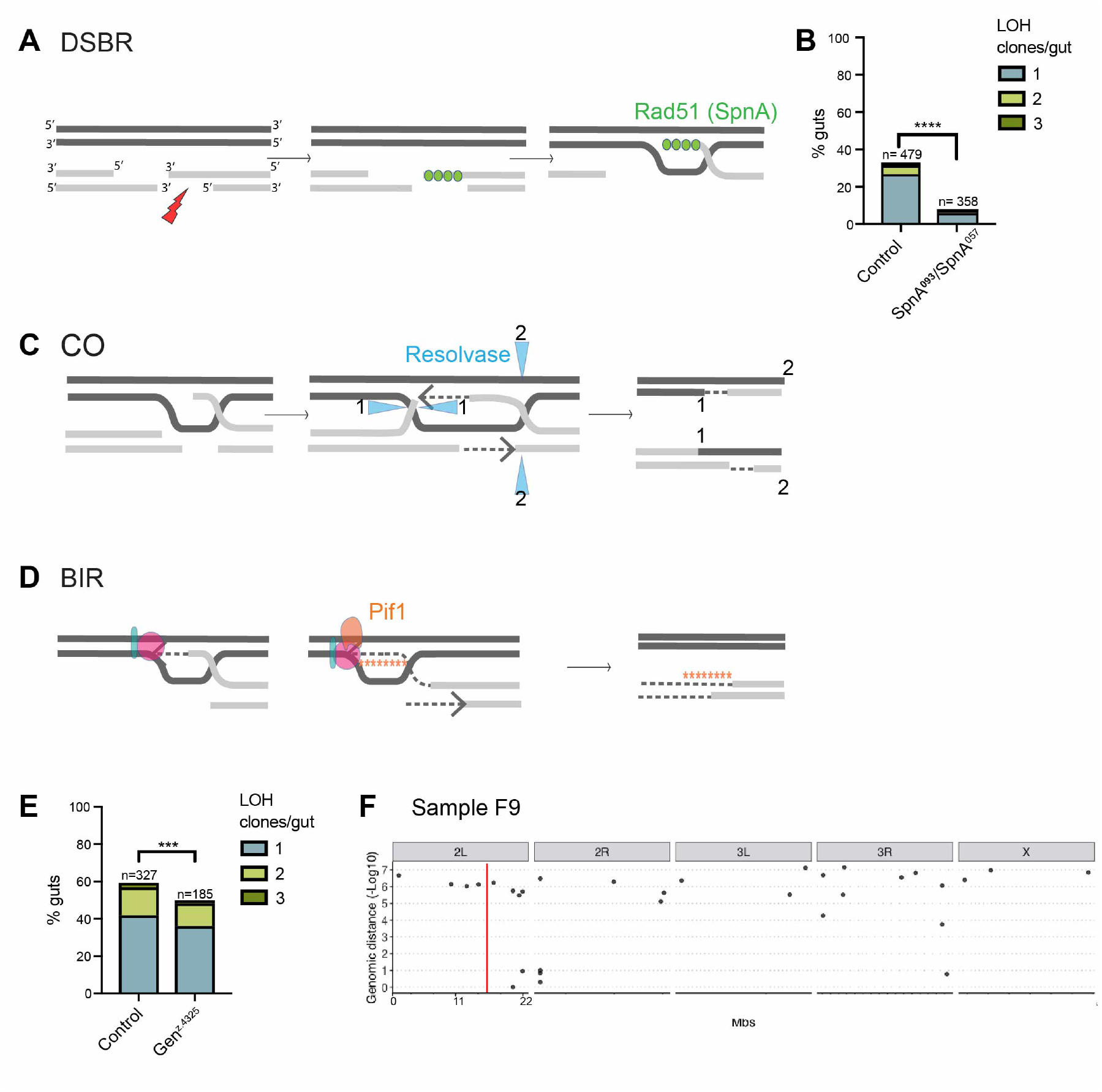
LOH largely arises from a Rad-51 dependent mitotic recombination consistent with double Holliday junction resolution. **(A)** Double-strand break repair (DSBR) is initiated by 5’ end resection and Rad51(SpnA) coating of a 3’ end. **(B)** Frequency of LOH events in control compared to SpnA null mutant. Fisher’s exact test p<0.0001. **(C)** A crossover (CO) mechanism is mediated by a dHJ and relies on resolvases. Cleavage at sites 1 and 2 result in CO occurring between homologous chromosomes. Depending on the DNA strand, the resulting repaired chromosome can either contain one long LOH tract, or intervening regions of gene conversion nearby the initial breakpoint. **(D)** Break-induced replication (BIR) relies on a Pif1 helicase to unwind the template that is used for repair. Repair of the broken chromosome occurs by leading strand copying of the template, and subsequent lagging strand synthesis. It is prone to mutagenesis (stars). The resulting chromosomes are shown, where one is unaltered and one has been repaired by the de novo synthesis. **(E)** Frequency of LOH events in control compared to gen null mutant. Fisher’s exact test p<0.0001. **(F)** Representative rainfall plot showing no mutation pileup by the mapped recombination site in sample F9. The red line indicates the site of mitotic recombination.

### Genetic requirements and molecular signatures support double Holliday junction resolution leading to crossing-over

Two distinct mechanisms of mitotic recombination could explain the long tracts of LOH that were detected in intestinal stem cells: (1) Crossover, resulting from a classic double-strand break repair involving a dHJ structure, whose resolution would lead to reciprocal exchange of segments of the homologous chromosomes **(Figure 6C)**. (2) BIR, in which an error-prone polymerase would copy material directly from the homologous chromosome **(Figure 6D)**. While both mechanisms could lead to long stretches of LOH spanning the chromosome arm and depend on *Rad51*, they differ in their genetic requirements. For this reason, we first decided to test roles of DNA repair proteins specific to each mechanism.

The resolution of dHJ DNA structures relies on resolvases such as Gen1 and Mus81 [34, 68]. While *mus81^nhe1^*mutant flies did not show consistent differences between mutant and control (**Supplementary Figure 6A),** the loss-of-function of Gen1 resolvase, encoded by *gen*, on the other hand, showed a significant decrease in LOH events reducing LOH from 59.3% in controls to 42.16% in *Gen1* mutants (**Figure 6E**). These data suggest that *Gen1* is important to promote LOH likely through facilitating DNA repair via dHJ resolution that results in crossover.

BIR relies on the DNA helicase Pif1, important to unwind the double-strand DNA thereby allowing to the copying DNA from the homologous donor chromosome [69–72] (**Figure 6D**). *Pif1* null mutant flies are not viable [70]. Therefore, *Pif1* RNAi was expressed specifically in adult ISCs though it did not significantly alter LOH frequency in *Su(H)-/+* flies (**Supplementary Figure 6B**), suggesting that Pif1 may not play an integral role in the repair giving rise to the LOH events, though inefficient knockdown cannot be excluded. We, therefore, also examined the genomic features of the chromosomes with LOH.

Taking advantage of the high density of parental SNPs, we could assess the regions surrounding the recombination site for evidence for potential molecular features associated with dHJ resolution or BIR. 16/20 samples showed a simple shift from heterozygous to homozygous, and appear to be homozygous throughout the chromosome arm (**Figure 4C**, **Supplementary Table 2**). In contrast, 4/20 samples had evidence for intervening tracts where SNPs would alternate from heterozygous to homozygous and then homozygous to heterozygous again, followed by heterozygous to homozygous and crossover (**Supplementary Figure 7A-C)**. For example, sample F1 had an initial shift from heterozygous to homozygous SNPs (at position Chr 2L: 20701878-20702379), then 1.5kb more distally, a region of heterozygosity was again detected, clearly supported by 2 informative SNPs, followed by again a shift from heterozygous to homozygous throughout the chromosome arm **(Supplementary Figure 7A)**. Similar shifts from heterozygous to homozygous and homozygous to heterozygous could be detected on 3 other samples (**Supplementary Figure 7B**, **7C,** and data not shown for sample F13 where more than 3 shifts were detected suggesting a complex event). These types of tracts could result from predicted outcomes of resolution of dHJ structures, mismatch repair of heteroduplex DNA arising from dHJ structures, or from template-switching previously shown to occur in 20% of BIR events [25, 73].

In order to further distinguish between mitotic recombination driven by dHJ resolution versus by BIR, we next assessed the accumulation of somatic point mutations, a hallmark of BIR but not dHJ processing ([74] and reviewed in [28]). Somatic point mutation (single-nucleotide variant; SNV) calling was performed on the genomic samples and none showed evidence of increased mutations near the recombination sites or throughout the arm of Chr 2L (**Figure 6F, Supplementary Figure 6C**). Therefore, these data argue against BIR as a primary mechanism driving LOH in ISCs.

Altogether, these data support and important role of LOH events being driven by cross-over arising from dHJ resolution, and not BIR. While this mechanism allows repair of double-strand breaks in stem cells promoting their survival, it leads to LOH of the tumor suppressor *Su(H)* and drives neoplasia initiation.

## Discussion

Our findings reveal essential intrinsic and extrinsic factors acting on DNA damage and repair in adult stem cells influencing LOH, an important mechanism of tumor-suppressor inactivation. Using whole-genome sequencing and genetic approaches, our data suggest that a major contributor to genome alteration in *Drosophil*a ISCs involves mitotic recombination-driven LOH that primarily occurs through a classic HR mechanism depending on dHJ resolution.

We find that differences in the intestinal microbiota can have distinct impacts on the mutation rate of a tissue. Infection by *Ecc15*, but not *Pe*, altered LOH frequency. *Ecc15* and *Pe* both induce damage to the intestinal epithelium and promote ISC proliferation [42, 43]. Interestingly, we find here that gut enteric pathogenic bacteria can also increase marks of DNA damage. In our experiments, *Ecc15* led to a higher amount of DNA damage than *Pe*, which could explain why *Ecc15* increased LOH frequency but not *Pe*. We also found that under the doses of bacteria that we used, *Ecc15* led to a more pronounced increase in proliferation that *Pe* as assayed 24 hours after infection. Of note, the effects of *Ecc15* on DNA damage and LOH can be explained by the increased amount of replication-associated DNA damage as expression of *CycE* and *Stg* in ISCs can also increase LOH frequency and DNA damage. It is possible, however, that other differences between these bacteria or their interaction with the midgut epithelium could also underlie the effect on LOH. Indeed, these bacteria induce overlapping as well as distinct changes in gene expression in the midgut [75]. In addition, at high doses, *P.e.* can inhibit the translation machinery and block ISC proliferation. It is possible that roles in translation or subtle differences in stem cell viability may preclude detections of changes in LOH frequency in *Pe* conditions. In humans, inflammatory bowel disease and colorectal cancer have been linked to alteration of the microbiome [76–78]. In particular, the NC101 strain of *Escherichia coli* causes DNA damage [79], and creates a specific mutational signature [80]. Determining how additional pathogenic bacteria and other changes to environmental conditions affect LOH events may provide strategies for cancer reduction or prevention.

Using high-resolution SNP-based mapping of recombination regions, our study provides direct evidence of mitotic recombination as an underlying mechanism occurring in adult somatic stem cells. Our genetic experiments support these molecular findings as we found reduced LOH in mutant contexts of *Rad51* (*spnA*), and *gen*. Our recent studies illustrated that, in addition to genome alteration by recombination described here, point mutations, deletions, complex structural variants, and transposable element mobility all alter the somatic genome of ISCs and could, in theory, contribute to LOH [59, 60]. Indeed, we did find one example of LOH driven by a large 1.38 Mb deletion event. However, our data strongly support mitotic recombination being the primary mechanism by which LOH is generated in ISCs. Importantly, we found no evidence supporting a BIR mechanism as no increase in point mutations was found at LOH regions for the 23 samples that we assessed as would be predicted from previous work [74]. While our study could only assess those events inducing LOH, it is likely that a much higher number of DNA double strand breaks are produced but repaired using the sister chromosome, which would be undetectable in our assay. Our findings that dHJ resolution is an important contributor to LOH in somatic tissues suggest that targeting this pathway may be beneficial in patients with germline mutations in recessive tumor suppressor genes or dominant disease genes.

One important question is whether the sites at which recombination arises are simply sites of DNA damage or whether certain underlying genomic features specifically promote crossover events. Previous studies mapping positions of crossover, for the most part, have been performed using low-resolution techniques that precluded fine mapping of DNA sequences. To our knowledge, our study is the first high-resolution mapping in a Metazoan somatic tissue. In a study of cell lines derived from colorectal cancer driven by LOH of APC, sites of mitotic recombination were resolved to low resolution (∼4.5 Mb) and the authors suggest that the sites were non-random and possibly associated with low copy repeats (LCRs) [16]. Previous findings mapped mitotic recombination within the *Drosophila* male germline at low resolution using marker genes concluded that sites were non-random and provided indirect evidence that they may be linked to problems arising during replication [33]. A more precise mapping has been achieved in yeast [81–84]. These studies found a mitotic recombination hotspot near sites of replication termination, inverted repeats of Ty transposable elements, and G-rich quadruplex sequences [81–84]. While our sample size (n=23) is limited, we did not detect enrichment at mapped crossover points of these sequences. However, we did find a significant enrichment over what would be predicted by chance at SIRS and the *Histone Locus-Cluster*. Our findings of enrichment of SIRS at recombination sites are consistent with previous reports of their ability to stall replication forks and their enrichmenr at translocation break-points in human cancers [85]. The *Histone Locus-Cluster* encodes a ∼100 tandem repeats of 5 histone genes (*Histone 1*, *2A*, *2B*, *3A* and *3B*). Why this region has an elevated number of recombination sites than would be predicted by chance is not currently clear, however, we speculate that it is related to the unique features of these genomic locus. One peculiarity of these genes (excluding *Histone H1*) is their exclusive expression at high levels specifically during S phase [67], at which time collisions with DNA polymerase could occur. In addition, this genomic region is also unique in that is forms a nuclear subdomain, the Histone-locus body, where specific *Histone* 3’ RNA processing occurs, as these are unique in lacking canonical poly A tails. Consistent with DNA damage at the *Histone Locus Cluster*, the *HIST1H* cluster in human B-cells is found to acquire marks of γH2AX [86]. The potential of other genes highly expressed during S phase to be hotspots of DNA damage merits further investigation.

Our study underscores the notion that adult organs are genetic collages, where somatic mutations in adult progenitor cells drive divergent genomes within a common tissue. Given the number of events that we detect here for a marker at one genomic location on 1 out of the 5 major chromosome arms, we can estimate that ∼40 LOH events occur per gut for distal genes near the telomeres. As there are ∼1000 ISCs per midgut, we can estimate that 1 in 25 ISCs has an LOH event, raising the likelihood that this mechanism of somatic genetic diversity could alter tissue dynamics during the ageing process. Aside from arising somatically in Metazoa, mitotic recombination also occurs in plants as well as diatoms, where it is proposed to contribute to adaptive evolution of clonal populations [87, 88]. Future studies will likely reveal both common as well as species-specific molecular and environmental mediators of mitotic recombination.

## Methods

### Drosophila stocks

The following fly stocks and alleles were used in this study: From the Bloomington stock center, P(UAS-GFP.nls) (BL 4776). The following stocks were generous gifts: *SpnA^093^, SpnA^057^, w^1118^* (M. McVey), *Gen1*Z^.4325^*, Mus81^nhe1^* (J. Sekelsky), *ProsV1Gal4* (J. de Navascués), *Su(H)^Δ47^*, (F. Schweisguth), *O-fut1^4R6^*(K. Irvine), *Dl^Gal4^* (S. Hou), *N*^55e11^ (F. Schweisguth), *UAS-DAP, UAS-CycE, Stg, tubGAL80ts; Dl-GAL4* (B. Edgar). From the Vienna *Drosophila* Resource Centre (VDRC): *Rad51(spnA) RNAi* (VDRC 13362), *Mus81 RNAi* (VDRC 33688), *Pif1 RNAi;*-(VDRC 34533). For all experiments of LOH, the second chromosomes assayed for LOH were identical between the controls and experimental samples.

### Drosophila husbandry

For standard aging of experiments in **Figures 1, 2A-E, 2I-L, 6B**, **6E, Supplementary Figures 5A-B, 6A,** female flies were maintained at 25°C on a standard medium composition. For aging experiments, adult progeny were collected at 25°C over 3-4 days. Females were aged with males in the same cage (plastic cages 1 cm diameter, 942 ml). 400-600 flies/cage. Freshly yeasted food was provided in petri dishes every 1-2 days. Flies were transferred every 7 days without CO_2_ anaesthesia to clean cages. Dead flies were scored upon each food change to assess survival rates. Female flies were dissected at 5-6 weeks unless otherwise indicated.

For experiments in **Figures 2F-H**, **Supplementary Figures 1B-C’**, **5C, 6B** the genetic background *Su(H)^Δ47^/ UAS-GFP*; *Dl^Gal4^ tub-GAL80ts*/ + was used to drive stem cell specific expression in the ISC, due to the *Dl^Gal4^* driver. Crosses were maintained at 18°C on standard medium, flies were also maintained at 18°C during development and metamorphosis. Newly eclosed flies were collected over 5-7 days. Flies were maintained at 29°C thereafter. The shift to 29°C induced ISC specific UAS-driven transgene as the Gal80 repressor is temperature sensitive. Flies were dissected after 1 week for *UAS-DAP, UAS-CycE, stg* or after 2 weeks at 29°C for *Rad51RNAi, Mus81 RNAi, Pif1 RNAi*.

### Bacteria Treatments

Adult *Su(H)^Δ47^/+* flies were treated for 24 hours on filter paper soaked with *Ecc15* / *Pe* or control solution (see below) covering sugar agar (1.5%) plates. *Ecc15*/ *Pe* treatment: a 1:1 mix of OD 200 *Ecc15* culture and 5% sucrose. Control: a 1:1 mix of LB and 5% sucrose. Treatment was repeated once per week for 3 weeks, followed by a 1-week-recovery before dissection at 5 weeks. Proliferation response was assayed by phospho-histone 3 staining 24 hours after treatment in young 3-5 day-old flies.

### X-ray induction

1 week old flies were placed in the X-ray generator CIXD and exposed to 40 Gray at the RadeXp facility at Institut Curie.

### Immunofluorescence

Midgut fixation and immunofluorescence staining were performed as described previously described in [61]. Adult female midguts were dissected in PBS and then fixed at room temperature (RT) for 2 hours in 4% paraformaldehyde. Guts were trimmed and incubated in PBS 50% glycerol for 30 minutes before equilibration in PBS 0.1% Triton X-100 (PBT) to clean the lumen. Fixed and cleaned guts were then washed in PBT for at least 30 min before addition of primary antibodies (overnight at 4°C or 3-5 hours at RT). After at least 30 min wash, secondary antibodies were incubated 3-5 hours before DAPI staining (1mg/ml) and mounted in 4% N-propyl-galate, 80% glycerol.

The following primary antibodies were used: mouse anti-Delta extra-cellular domain (1/1000; DSHB C594.9b 10X concentrate), mouse anti-Pros (1/1000; DSHB MR1A-c 10X concentrate); chicken anti-GFP (1/2000, Invitrogen #A10262), rabbit anti-PH3, (1:1000; Millipore #06-570 lot 3746384), mouse anti-γH2Av (1:100; DSHB #UNC93-5.2.1 10X concentrate).

Imaging was performed using Zeiss LSM900 and LSM780 confocal microscopes and epifluorescence widefield microscope at the Curie Institute imaging facility with serial optical sections taken at 1 μm intervals (512X512 or 1024X1024) using 20X or 40X oil objectives through the whole-mounted posterior midguts.

### Quantification

Quantification was carried out blind. Results quantified are pooled data from either 3 biological replicates in **Figures 2L**, **6E** and **Supplementary Figure 5C, 6A**, **6B** or pooled data of 4 biological replicates in **Figures 2E**, **2G, 6B**.

#### LOH scoring

LOH events in females heterozygous for *Su(H)* were scored as clusters of at least 20 Delta and/or Prospero positive diploid cells.

#### Scoring of *Su(H)* LOH clones in Dl-Gal4 background using UAS-GFP

Experiments in **Figures 2G**, **Supplementary Figures 1B-C’**, **5C, 6B** the genetic background *Su(H)^Δ47^*/ *UAS-GFP*; *Dl^Gal4^*/ + was used to drive stem cell specific expression in the ISC, due to the *Dl^Gal4^*driver. However, as *Dl^Gal4^* is a loss of function allele of *Dl*, also a *Notch* signaling component, two types of neoplastic LOH clone could arise: those where *Su(H)* is inactivated and those where *Dl* undergoes LOH. We distinguished between these two possibilities by taking advantage of a UAS-GFP transgene located on Ch2L more distal on the chromosome arm to *Su(H)*. Therefore, LOH through recombination of *Su(H)* results in neoplastic clones that are GFP negative. These events were scored. In contrast, LOH resulting from other events including mitotic recombination of 3R leading to Dl LOH, are GFP+. These events were scored, but not counted in the analysis. Statistical analysis for LOH clones was carried in out in Prism using Fisher’s exact test (two-tailed) was performed and significant values were reported as: * p<0.05, *** p<0.001; **** p<0.0001, ns=not significant.

#### PH3 quantification

PH3 positive cells in the midgut were counted using the epifluorescence microscope.

#### γH2Av quantification

Images were acquired on the LSM900 confocal microscope. A maximum Z-projection was generated for all images on Image J (FIJI version 1.0). Only nuclear γH2Av intensity in the ISCs was measured.

### Sample collection for whole-genome sequencing

*Su(H)^Δ47^/+; ProsV1Gal4; UAS-nlsGFP* were used to visually identify midguts containing LOH neoplasias. The region of the LOH neoplasia was manually microdissected. An estimate of 50-80% purity of the LOH neoplasia can be achieved. These neoplastic LOH tumors were dissected together with the fly head. Genomic DNA was isolated using QIAamp DNA MicroKit (Qiagen) according to the manufacturer’s instructions.

#### Library preparation

Library preparation was performed with the Nextera XT kit and KAPA Hyper Plus ROCHE kit by the NGS facility of the Institut Curie. Samples were sequenced on one full flow cell (1600M clusters) on the NovaSeq in a paired-end 150bp mode.

Nextera XT Sample Prep Kit (Illumina) was used to prepare DNA sequencing libraries from 0.5 ng of genomics DNA. A step of enzymatic tagmentation with Nextera transposome was done to fragment DNA and add adapter sequences (Unique Dual Indexing strategy). A final amplification of the library was then performed with 12 cycles. After qPCR quantification (KAPA library quantification Kit, Roche), sequencing were performed on the Illumina NovaSeq 6000 using 2 x 150 cycles to get ∼40 M paired-end reads per sample.

KAPA Hyper Plus Kits (Roche) was used to prepare DNA sequencing libraries from 1.3 ng of genomics DNA. A first step of enzymatic fragmentation of 20 minutes at 37°C was done. An end-repair and A-tailing step on dsDNA fragments have produced end-repaired dsDNA fragments. A ligation overnight with the adapters (Unique Dual Indexing strategy) and a final amplification of the library was then performed with 15 cycles. After qPCR quantification (KAPA library quantification Kit, Roche), sequencing were performed on the Illumina NovaSeq 6000 using 2 x 150 cycles to get ∼40 M paired-end reads per sample.

### Bioinformatics

#### Analysis of whole-genome sequencing data

Sequencing analysis of LOH samples was performed using custom scripts implemented in the Nextflow (Di Tommaso, 2017) pipeline nf-lohcator https://github.com/nriddiford/nf-lohcator.

Briefly, adapter sequences were removed using trimmomatic 0.39 [91]. Trimmed reads were aligned to release 6.12 of the *Drosophila* melanogaster reference genome (FlyBase) using bwa-mem version 0.7.17 (Li H, 2013, arXiv:1303.3997v2 [q-bio.GN]). Fastqc (Andrews, D. (2010)) FastQC: A Quality Control Tool for High Throughput Sequence Data [Online]. Available online at: http://www.bioinformatics.babraham.ac.uk/projects/fastqc, bamstats (https://biopet.github.io/bamstats/1.0.1/index.html) and multiqc [92] were used to provide quality control files.

Alignment files were sorted and indexed using SAMtools [93] followed by the generation of pileup output summarizing the base calls of aligned reads to the reference sequence using mpileup command in SAMtools for each sample. VarScan2 version 2.4 (https://github.com/dkoboldt/varscan/releases) was then run on pileups generated. “Lohcator”, a custom python script https://github.com/nriddiford/nf-lohcator/blob/master/bin/lohcator.py developed in the lab was then ran to identify shifts in zygosity for each tumor sample and to create bed files to facilitate identification of the start and end of LOH for each sample.

#### VAF representation

VAF plots in **Figure 4B**, **Supplementary Figures 1A, 2A, 3A, 3B** were plotted with alleleFreqs: https://github.com/nriddiford/alleleFreqs using the LOH calls from VarScan2.

#### Rainfall plots

To generate rainfall plots in **Figure 6F** and **Supplementary Figure 6C**, filtering was carried out by using the “high quality” somatic calls produced by VarScan2.

We specified minimum coverage of 20 for both the tumour and normal sample and a somatic p-value<0.05. These calls were represented on rainfall plots. Some samples showed no SNVs and are thus not represented in the **Supplementary Figure 6C**.

#### Mapping of regions of recombination

First, on IGV, we confirmed that at this region, all the informative SNPs in the head are heterozygous and have gone homozygous in the tumour. Using a tool we have developed to identify regions of LOH in matched tumour normal pairs (nf-lohcator, see above), we were able to determine where the first homozygous SNP (from the centromere) is located, we then manually located the first heterozygous SNP on IGV relative to that for the mitotic recombination events. It is important to note that both coverage and tumour purity play an important role in confidently mapping these regions, with more emphasis placed on purity. We determined values for both coverage and purity for each mapped region (see **Supplementary Table 1**). We were unable to map the region of recombination for 3 samples (Samples F8, F10 and F26) because of EC contamination. These samples however allowed us to benchmark what is deemed “too impure” for the mapping of regions of recombination.

#### Association of mapped breakpoints with genomic regions

To assess whether mapped breakpoints were enriched for *The Histone Locus Cluster* **(Figure 5B)**, and other genomic features **(Supplementary Figure 4A-D)**, permutation tests were performed using regioneR [94] to determine the significance of the overlap between sequence features and our mapped breakpoint regions. In order to compare observed counts between real and shuffled data, we restricted permutations to within the genomic locus where mitotic recombination can be detected between the *Su(H)* locus and the centromere chr2L:15039488-23512838, and performed 10,000 permutations.

#### Calculating coverage

Mosdepth was used to calculate genome-wide sequencing coverage with default parameters (https://github.com/brentp/mosdepth).

#### Copy number analysis

Copy number analysis was performed using a read-depth-based approach with CNV-seq [95]. CNVPlotteR https://github.com/nriddiford/cnvPlotteR.git was then used to generate copy number plots shown in **Supplementary Figure 2B** and **Supplementary Figure 3B’**.

#### Calculating EC contamination

The approach described in [60] and implemented in https://github.com/nriddiford/winlow was used to determine the likely contamination of tumor samples with EC cells (**Supplemenary Table 2**).

#### Data Availability

The data have been deposited with links to BioProject accession number PRJNA858414 in the NCBI BioProject database (https://www.ncbi.nlm.nih.gov/bioproject/).

## Acknowledgements

The authors would like to thank members of the Bardin team, Sarah Lambert, Yohanns Bellaïche, Pierre-Antoine Defossez, Evan Dewey, and Jeff Sekelsky for helpful discussions and comments on the manuscript. Primary grant support for this work was from Worldwide Cancer Research (WCR 20-0147 grant to A.J.B.). Additional financial support came by grants from Fondation pour la Recherche Médicale (A.B., DEQ20160334928), as well as funding from the program “Investissements d’Avenir” launched by the French Government and implemented by ANR, ANR SoMuSeq-STEM (A.B and N.S.), Labex DEEP (ANR-11-LBX-0044), IDEX PSL (ANR-10-IDEX-0001-02 PSL) and ICGex STEM-MITO-REC (A.B.). High-throughput sequencing has been performed by the ICGex NGS platform of the Institut Curie supported by the grants ANR-10-EQPX-03 (Equipex) and ANR-10-INBS-09-08 (France Génomique Consortium) from the Agence Nationale de la Recherche “Investissements d’Avenir” program), by the Canceropole Ile-de-France and by the SiRIC-Curie program–-SiRIC Grant “INCa-DGOS-465””. Salary support of A.J.B by CNRS; L.A-Z received funding from the European Union’s Horizon 2020 research and innovation program under the Marie Skłodowska-Curie grant agreement No 666003 and the French foundation for medical research (FRM, FDT201904008016).

## Author Contributions

L.A. A.J.B. designed the study and analyzed the data. N. Riddiford developed software and performed most of the bioinformatic analysis, with N. Rubanova. performing additional bioinformatic analyses supervised by N.S. L.A. and M.S. performed the experiments. M.B. from the ICGex NGS platform optimized low-input DNA sequencing. L.A. and A.J.B. wrote the manuscript with contribution from the other authors.

The authors declare no competing interest.

## Supplementary Figures

**Supplementary Figure 1:**
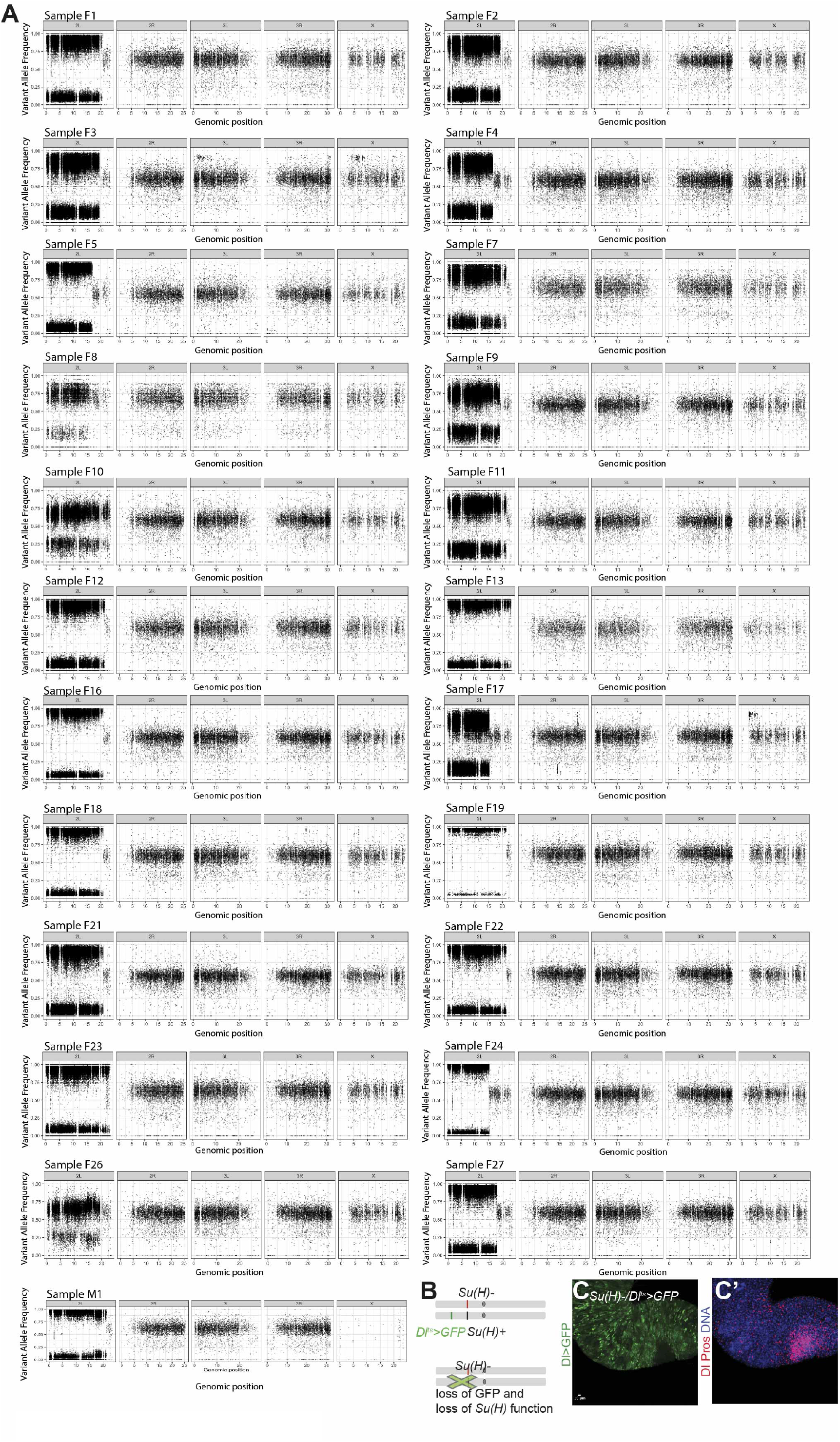
Evidence for LOH driven by Mitotic Recombination on chromosome 2L. **(A)** Variant allele frequency plots of samples supporting LOH of chromosome 2L, with SNPs becoming homozygous (VAF>0.75). Samples F6, F14, F15 and F25 were excluded because of too low coverage. **(B-C’)** The genetic background Su(H) ^Δ47^/ UAS-GFP; Dl^Gal4^/ + was used to drive stem cell specific expression in the ISC, due to the DlGal4 driver. Neoplastic LOH clones where Su(H) is inactivated resulted in GFP negative clones due to the loss of the UAS-GFP transgene located on Ch2L more distal on the chromosome arm to Su(H) via mitotic recombination (GFP in green, Dl and Pros in red, DAPI marking DNA in blue).

**Supplementary Figure 2:**
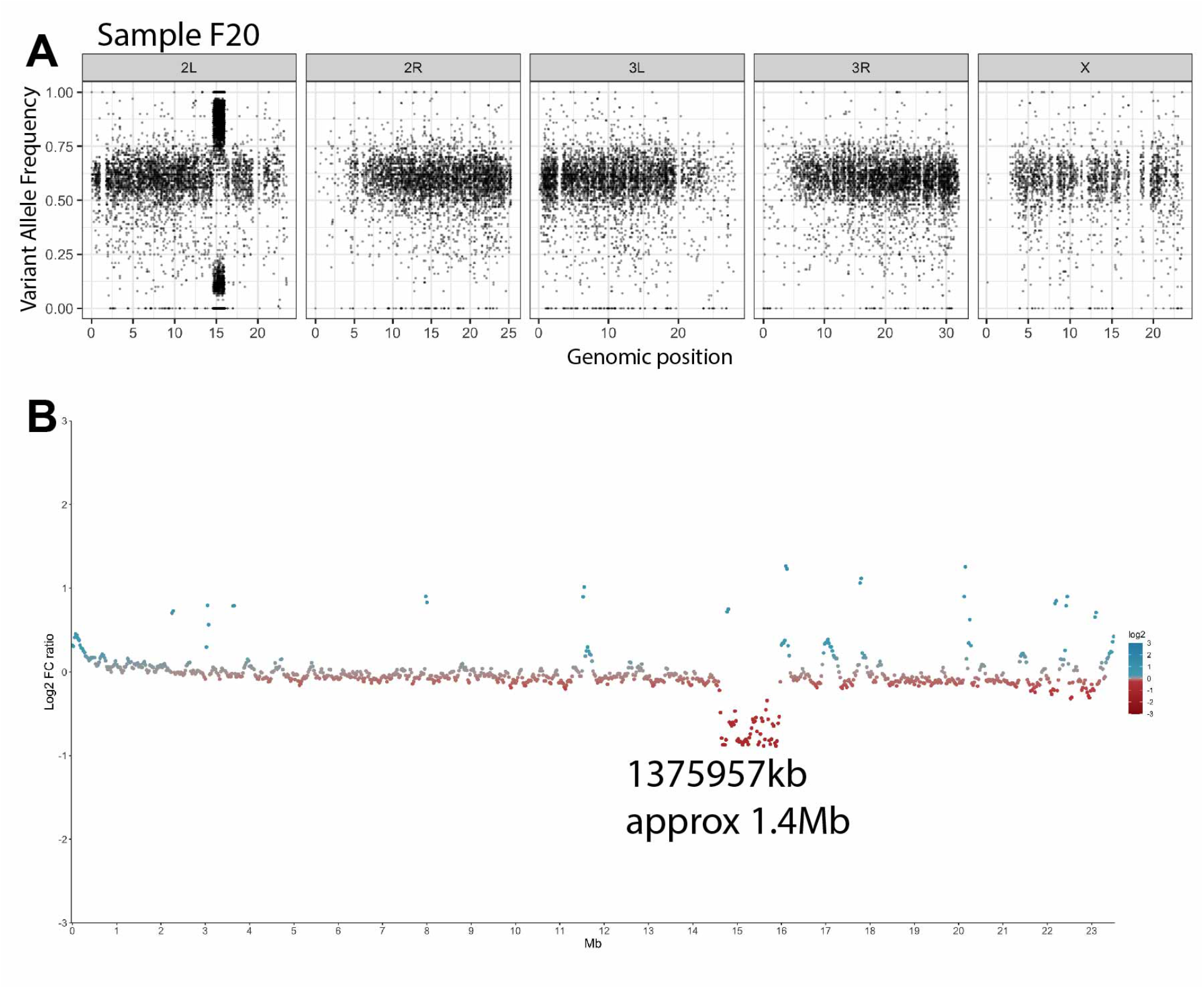
Sample showing a deletion spanning Su(H) locus leading to LOH. **(A)** VAF plot of sample F20 showing a shift in allele frequency on Chr 2L where there is a deletion. **(B)** Copy number plot showing a large drop in coverage (approx 1.4Mb) in the tumor relative to the head control.

**Supplementary Figure 3:**
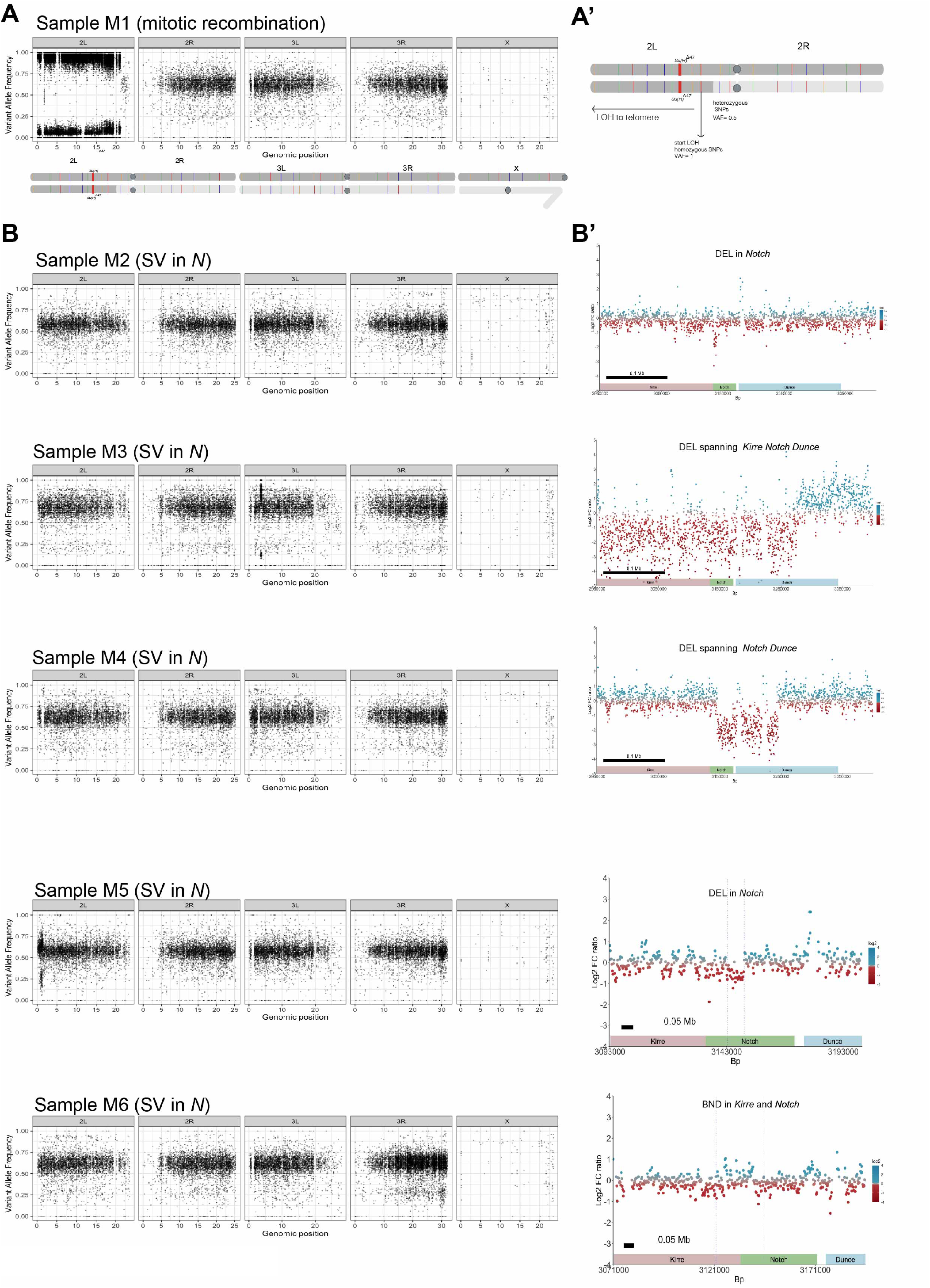
VAF plots of all the male samples. Plots of all male samples, showing all chromosomes. Only sample M1 **(A-A’)** shows that chromosome 2L has undergone an LOH event, with SNPs becoming homozygous (VAF >0.75). The remaining samples **(B-B’)** show no LOH on chromosome 2L (left panels) but rather show a structural variants (SV) spanning the Notch locus on the X chromosome (right panels). DEL is deletion, BND indicates “break-end class” of genomic aberration,.

**Supplementary Figure 4:**
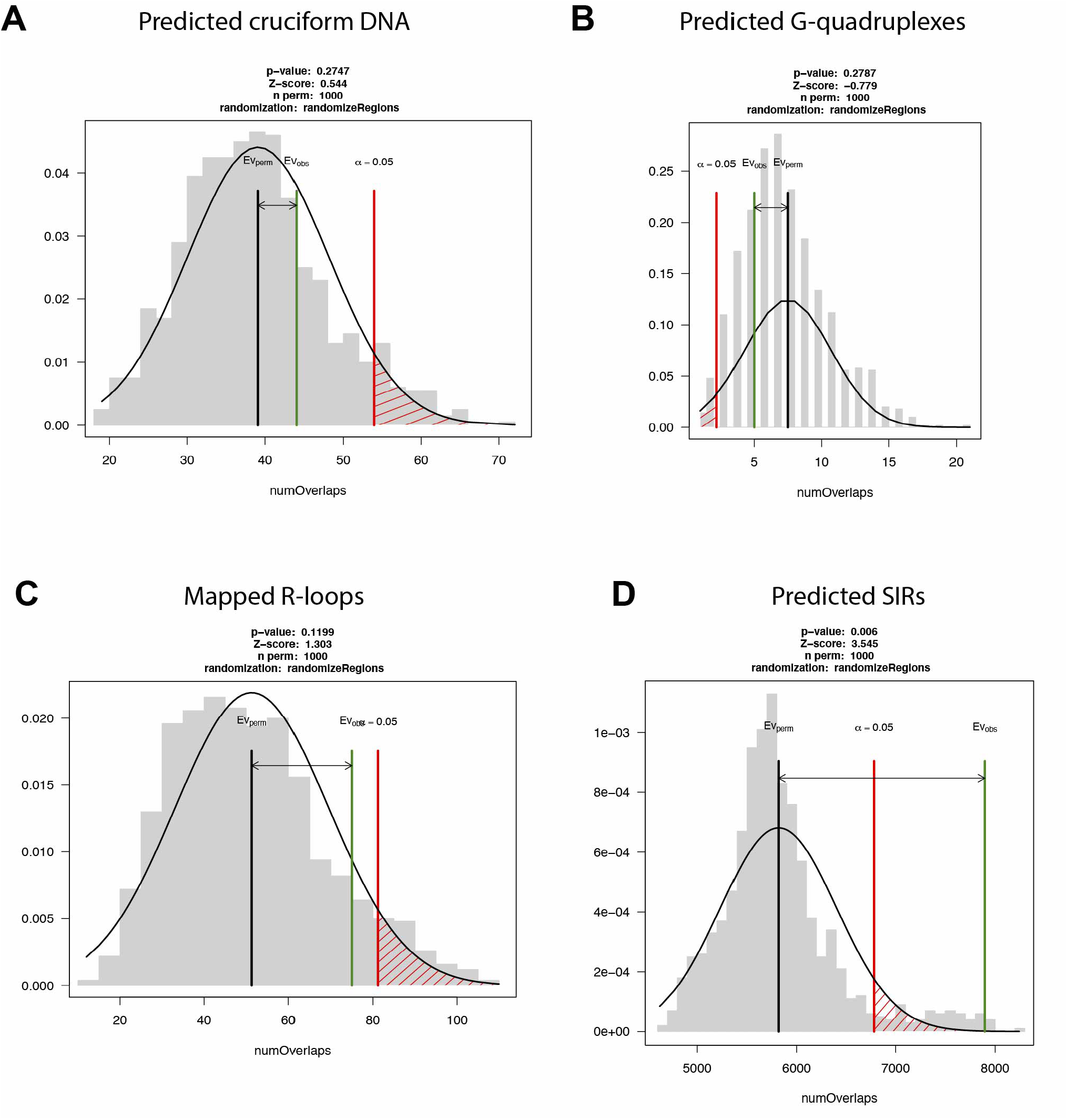
Mapped recombination sites showed no significant overlap with non-B-form DNA such as cruciform DNA, G-quadruplexes and R-loops. Permutation test carried our using RegioneR. Test association between mapped recombination sites and **(A)** predicted cruciform DNA [89], **(B)** predicted G-quadruplexes [90]. **(C)** mapped R-loops [66], **(D)** predicted SIRs [85]. Shuffles: 1000 constricted to the region between Su(H) and the centromere where recombination can be detected.

**Supplementary Figure 5:**
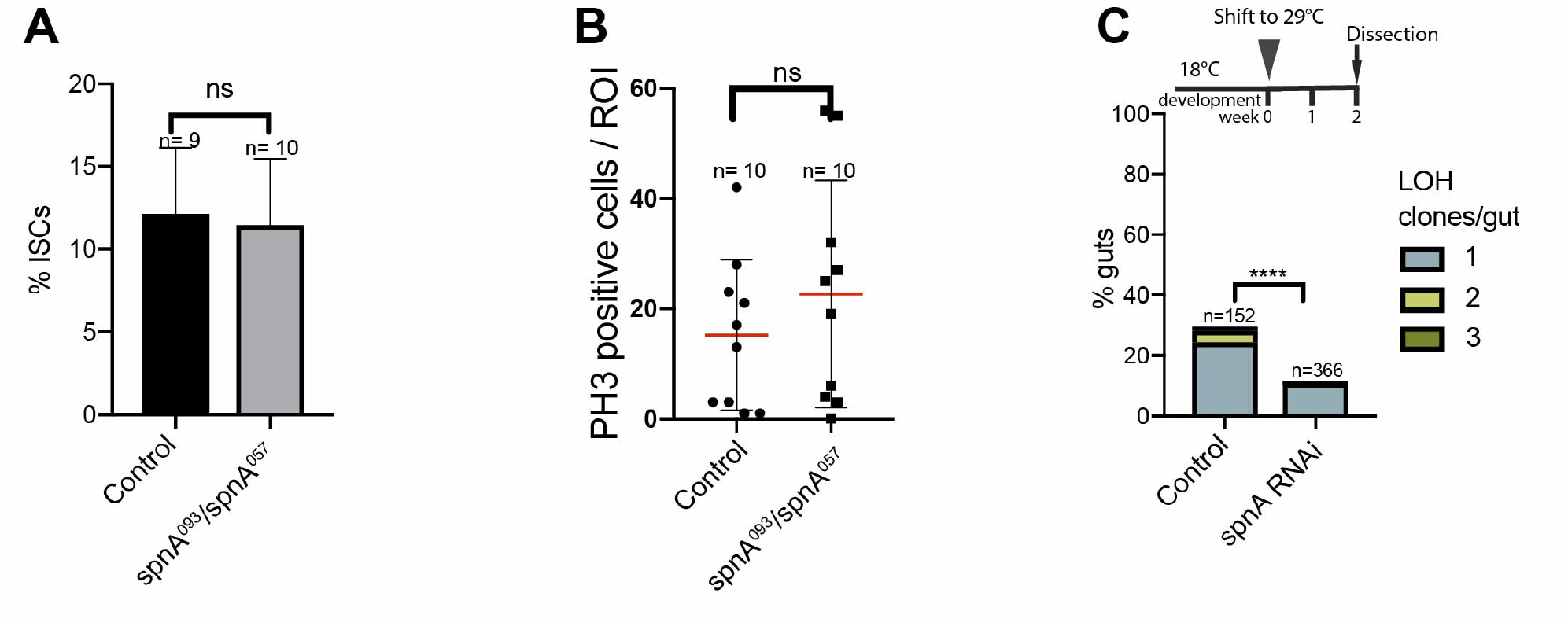
Evidence for SpnA promoting loss of heterozygosity. **(A)** No significant difference in %ISCs per region of interest (ROI) between control guts and Su(H)^Δ47^/+; SpnA^093^/SpnA^057^ (T-test with Welch’s correction). **(B)** No significant difference in mitotic cells per ROI between control guts and Su(H)^Δ47^/+; SpnA^093^/SpnA^057^ (T-test with Welch’s correction). **(C)** Frequency of LOH clones in control compared to Rad51 knockdown in ISCs. Adult flies were shifted to 29°Cto induce RNAi for 2 weeks then dissected. (Genotypes: SpnA RNAi; Su(H)^Δ47^/ UAS-GFP; Dl^Gal4^ tubGAL80^ts^ /+ Vs Su(H)^Δ47^/ UAS-GFP; Dl^Gal4^ tubGAL80^ts^ /+ control) (Fisher’s exact test, two-tailed).

**Supplementary Figure 6:**
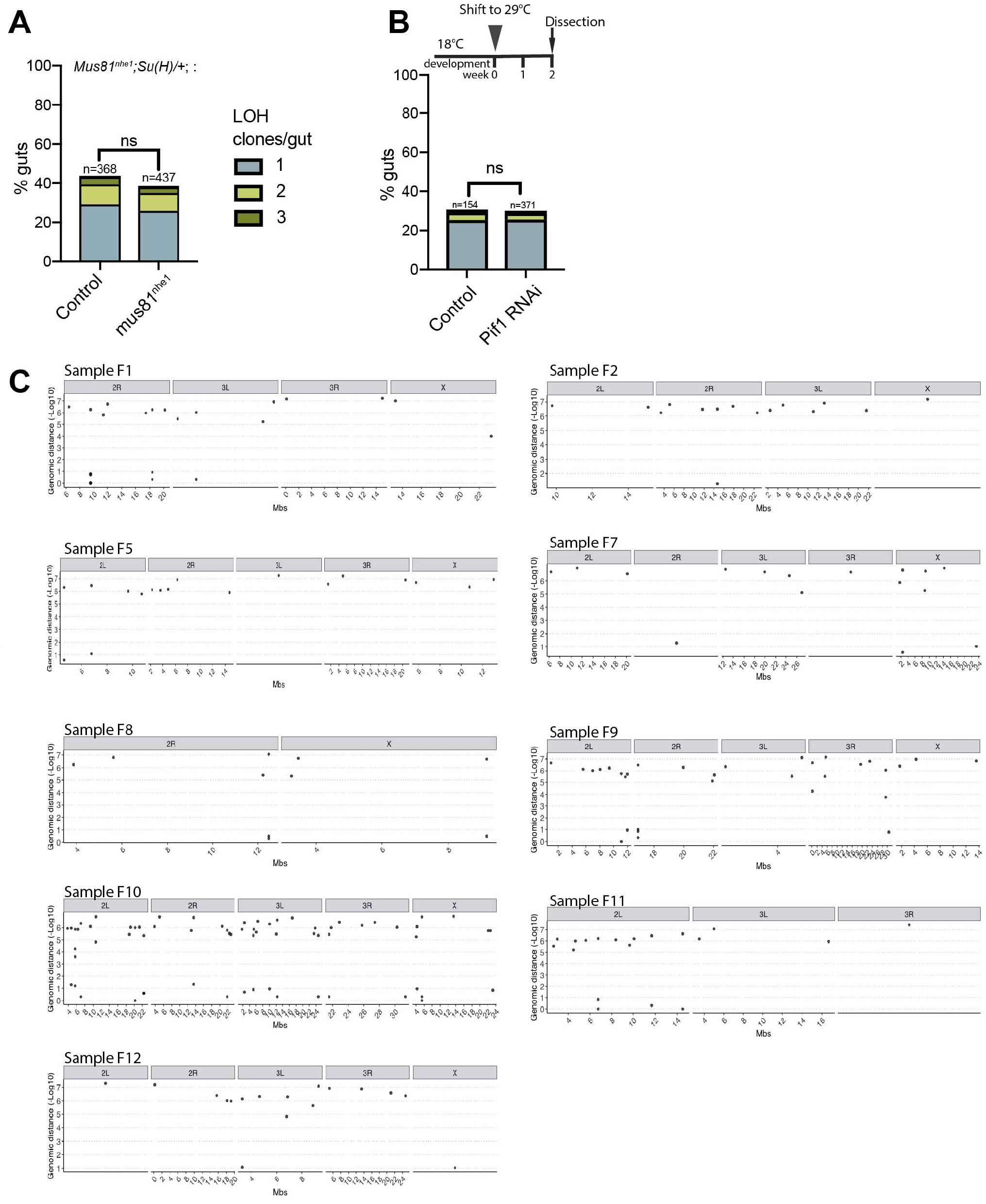
Genetic and genomic data arguing against a BIR model. **(A)** Frequency of LOH clones in Mus81^nhe1^; Su(H)/+ compared with +/+; Su(H)/+; control (Fisher’s exact test, two-tailed). **(B)** Frequency of LOH clones in control compared to Pif1 knockdown in ISCs. RNAi was induced in adult ISCs for 2 weeks prior to dissection. Genotypes : Su(H)^Δ47^/ UAS-GFP; Dl^Gal4^ tubGAL80^ts^ / UAS-Pif1 RNAi Vs Su(H)^Δ47^/ UAS-GFP; Dl^Gal4^ tubGAL80^ts^ /+ control. Fisher’s exact tests (ns). **(C)** Rainfall plots of point mutations in the sequenced LOH samples. Each point represents the genomic distance between two SNVs (point mutations). A mutational pileup would be detected by an accumulation of points vertically. We observed no mutational hotspots on chromosome 2L, arguing against a BIR model. Samples not shown are samples which had no SNVs.

**Supplementary Figure 7:**
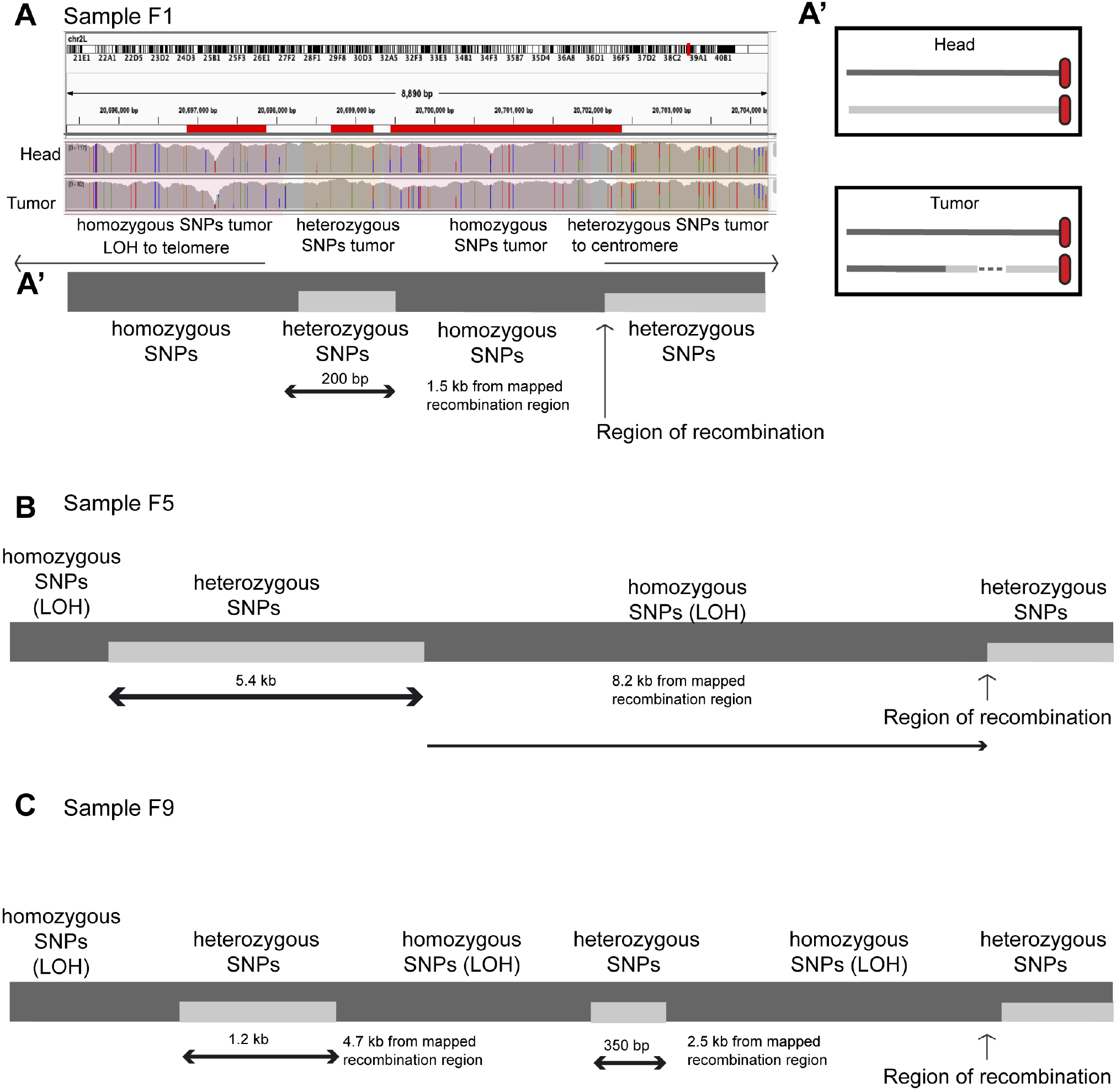
Schematic showing samples with conversion tracts. **(A)** IGV view showing SNP evidence of a conversion tract 1.5kb away from the mapped region of recombination. While the control head sample representing the germline (top IGV tract) shows all heterozygous SNPs, the bottom LOH tumor sample shows a shift from heterozygous→homozygous SNPs, then goes from homozygous→heterozygous, before going back from homozygous→heterozygous throughout the chromosome arm. A schematic of the cell of origin in head and tumor is shown on the right panel A’. **(A’)** Schematic of sample F1 conversion tract denoted by shift from homozygous to heterozygous SNPs in the tumour sample for ∼200 bp before again becoming homozygous until the telomere. **(B)** Schematic of sample F5: After the initial recombination site (indicated by arrow) there is a shift from heterozygous ->homozygous SNPs in the tumor, a region approximately 8.2kb away showed a DNA tract marked by a shift from homo-het SNPs for ∼5.4kb before again becoming homozygous until the telomere. **(C)** Schematic of sample F9 where 2 tracts were identified having heterozygous SNPs within the larger LOH region. Approximate length of each segmented is noted.

